# Modern Times: Longitudinal Study of Toba/Qom Communities Reveals Delay and Shortening of Sleep in Real-Time

**DOI:** 10.1101/2025.07.28.667301

**Authors:** Leandro P. Casiraghi, Ignacio Spiousas, Laura L. Trebucq, M. Florencia Coldeira, Camila R. Godoy Peirone, Victor Y. Zhang, Malen D. Moyano, Diego A. Golombek, Horacio O. de la Iglesia

**Author notes:** Corresponding authors: Leandro Casiraghi, Horacio de la Iglesia. CIESAS Sureste, San Cristóbal de las Casas, Chiapas, México (Camila R. Godoy Peirone).

## Abstract

While artificial light and digital technologies are widely assumed to delay and curtail sleep, direct, longitudinal evidence documenting these changes is remarkably scarce. From 2012 to 2024, we conducted a longitudinal study of native Toba/Qom communities in northern Argentina, including rural groups newly introduced to electricity and semi-urban groups with earlier access to electricity. Using linear mixed-effects models on a dataset of over 12,000 actigraphically recorded sleep events from 156 participants at different times, we observed striking shifts in sleep dynamics. Remarkably, both groups showed delays of up to 1.4 hours in sleep timing, and participants from rural communities lost a full hour of sleep within just a decade. These changes occurred across a period that included, first, the introduction of electricity to rural communities after 2016, and, in more recent years, the spread of the internet and smartphone use among the Toba/Qom. Our models reveal that these events alone cannot fully explain the observed shifts in sleep timing, suggesting that broader processes of modernization and acculturation, beyond electric light and digital devices alone, may be driving sleep changes. The rapid transformations observed in these communities mirror, within a single decade, the likely trajectory of human sleep across the 20th and early 21st centuries, underscoring the complex ways modern environments reshape sleep behavior.

## Introduction

The harnessing of electric power and thus the invention of the electric bulb revolutionized human life in ways that are impossible to quantify. One of its main effects was the conquest of the night hours, historically reserved for sleep, through the creation of lit environments safe from the perils of darkness. Electric light allowed for the extension of wakefulness and chores into the evening hours and, eventually, the birth of the 24/7 society of work and entertainment in the current hyper-industrialized world. This process has not only significantly delayed sleep times but has also likely driven a reduction in sleep duration. A poll by Gallup published in April 2024 reveals that Americans report they sleep markedly less than 80 years ago ^1^, and a systematic review of available children’s data from 1905 to 2008 suggests a decrease in daily sleep of one hour in this period ^2^. Nevertheless, whether sleep has shortened with industrialization remains controversial, and, indeed, a recent analysis of published sleep data suggests that the opposite may be true ^3^.

In order to study the ways in which industrialization has affected human behavior, we need to be able to study populations in which this process is still ongoing. Pre-industrial societies offer unique opportunities to investigate the impact of the adoption of modern technologies and their related habits on sleep patterns, as they usually display differential access to electricity, high levels of exposure to the natural environment, weaker scheduling impositions, and rely more on environmental and ecological time references ^4–7^.

Our group is focused on the study of native Western Toba/Qom populations in the northern region of Argentina ^8^. These people, formerly hunter-gatherers, remain a very isolated and underprivileged group spread in the Formosa province, relying mostly on government assistance for subsistence. For the last 14 years, we have been studying sleep and circadian rhythms of both rural and urban-adjacent Toba/Qom communities, allowing us to contrast the behavior of communities in which the most relevant distinguishing characteristic was, until not so long ago, their access to electricity. Our previous work has shown that, indeed, urban Toba/Qom displayed later sleep onsets and shorter sleep duration than rural communities with no access to electricity ^9^. This correlation between electric light and delayed sleep has also been reported by several other groups studying pre-industrial communities around the world ^10–13^.

Since we started working with Toba/Qom communities in 2012, electricity has been progressively introduced to most rural areas in this region of the country, including the most isolated settlements (Fig. S1). Since such efforts started around 2016, all the communities we have worked with are now connected to the electric grid, and at the time of writing this paper, every single family we have worked with has electricity in their homes. In later years, television sets, smartphones, and internet connections, while in limited supply, are already available to many families.

We have longitudinally monitored sleep patterns through wrist-actigraphy in these Toba/Qom communities at different seasons and time points since 2012, generating a dataset of over 12,000 sleep records. This dataset represents a unique opportunity to assess the impact of the introduction of electricity on the sleep of these communities in real-time. The study we present here provides a close insight into the history of human sleep patterns in developing pre-industrial communities, highlighting the profound impact of modern technology on human behavior.

## Results

Our sleep dataset from Toba/Qom communities includes over 12,000 sleep events recorded through wrist-actigraphy between 2012 and 2024 from a total of 156 participants (94 in the rural group, 62 in the urban group). Table S1 details the number of participants, recorded events, and age and sex composition of all data according to community, year, and dates when it was collected. Starting around the year 2016, electricity was progressively introduced in the rural communities in the Vaca Perdida and neighbouring areas including the communities we studied (Fig. 1.A and Fig. S1). Throughout this period of significant transformations for the rural Toba/Qom, we detect a progressive delay and shortening of sleep both at the individual and community level (Fig. 1.B).

**Fig. 1.**
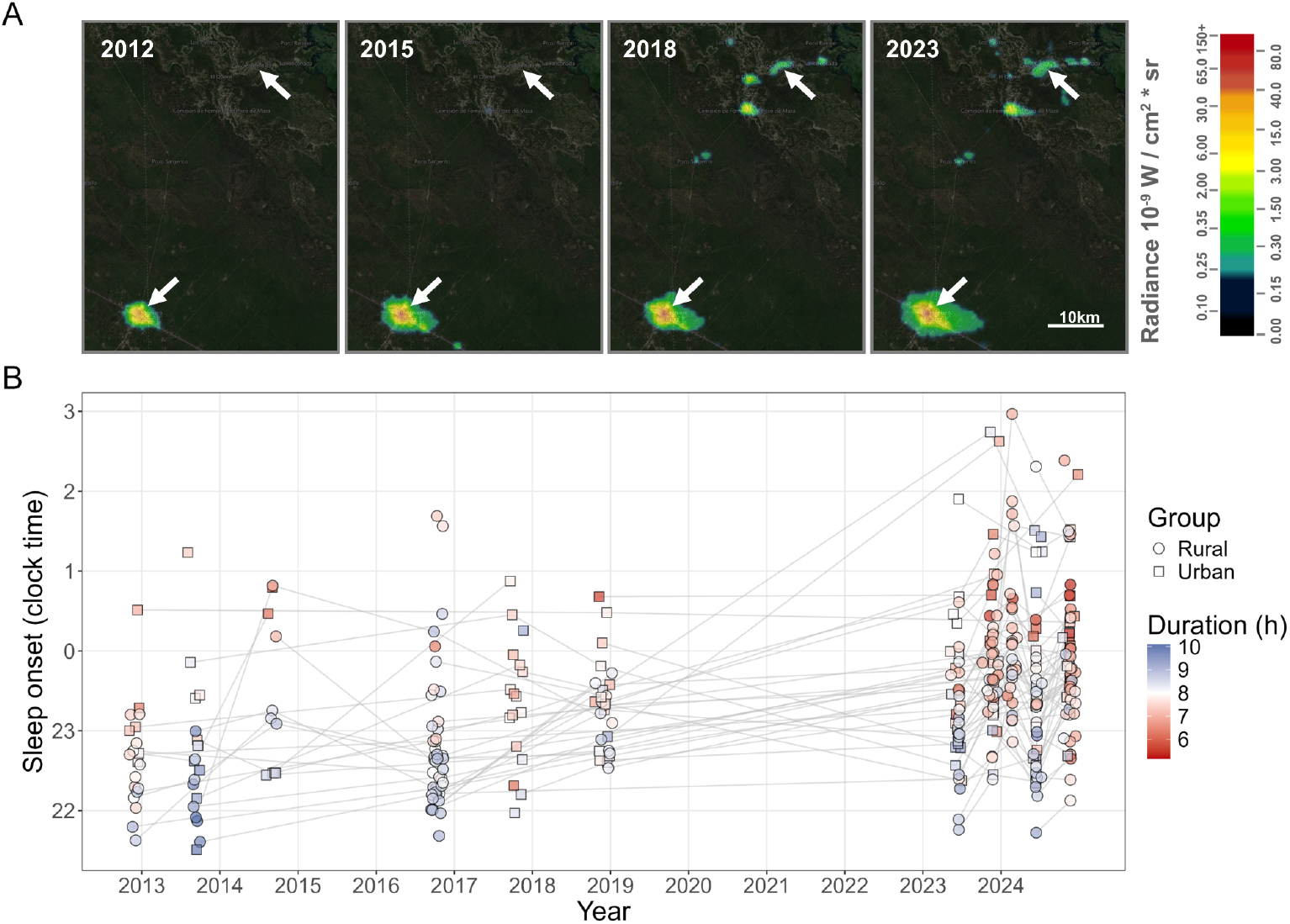
Evolution of night light pollution and sleep from 2012 to 2024 in both rural and urban areas. **A)** Radiance DNB levels measured by NASA’s VIIRS instrument in the geographic regions of Ingeniero Juárez (bottom-left arrow) and Vaca Perdida (top-right arrow) over the years, obtained from Jurij Stare’s Light Pollution Map. **B)** Summary of sleep start times and duration. Each marker indicates the mean time of sleep start for a participant at a single data collection campaign, with gray lines connecting repeated measurements. Fill colors represent the mean sleep duration, and shapes indicate the participant’s group.

Sleep start, midpoint, and end times, sleep duration, and sleep regularity (measured by the Sleep Regularity Index) were analyzed through linear mixed-effects models including groups (urban or rural), date, photoperiod length, type of day (workday or free-day), and sex and age as fixed effects, and including repeated measures per participant as a random intercept. Interaction terms between group and date, and between sex and age were also included in the models (see the Methods for full details on the model specifications).

Table 1 displays the full results of models for sleep onset, offset, and midpoint; Table 2 shows the results for sleep duration and sleep regularity. Coefficients are expressed along their 95% CI. The models revealed photoperiod length as a strong predictor of sleep times, with 0.18 [0.15-0.21] h delays in sleep start times (t = 12.235, p < 0.001), 0.17 [0.14-0.19] h advances in sleep end times (t = 12.444, p < 0.001), and a reduction of 0.35[0.32-0.38] h of sleep (t = 21.749, p < 0.001) per hour of increase in daylight. A longer photoperiod was also associated with slightly higher sleep regularity (1.2 [0.6-1.9] units/hr of photoperiod extension, t = 3.596, p < 0.001). Weekends displayed slightly later sleep start times (0.19 [0.13-0.24] h, t = 6.666, p < 0.001) and later sleep offset times (0.21 [0.16-0.26] h, t = 8.011, p < 0.001), and thus did not affect sleep duration. Accordingly, the midpoint of sleep was minimally delayed in nights before free days (0.2 [0.15-0.24] h, t = 9.014, p < 0.001), indicating that social jet lag is minimal in these communities. Male participants started their sleep significantly later (0.68 [0.41-0.95] h, t = 4.916, p 0.001), woke up later (0.56 [0.262-0.86] h, t = 3.689, p < 0.001), and had a lower sleep regularity (-3.3 [1.1-5.6] SRI units, t = 2.915, p = 0.004), but slept about the same amount as female participants.

**Table 1.** Linear models including random effects of sleep start, end, and midpoint times, expressed as hours from 00:00 on the date of start of sleep.

|  | Parameter | Coefficient | SE | 95% CI | t | p |
| --- | --- | --- | --- | --- | --- | --- |
| Sleep onset | (Intercept) | 22.95 (1) | 0.11 | [22.73, 23.17] | 207.673 | <0.001 |
|  | group = Urban | -0.10 | 0.12 | [-0.34, 0.15] | -0.787 | 0.431 |
|  | date (years) | 0.14 | 0.01 | [0.12, 0.16] | 12.630 | <0.001 |
|  | photop. length (h) | 0.18 | 0.01 | [0.15, 0.21] | 12.235 | <0.001 |
|  | day = weekday | -0.19 | 0.03 | [-0.24, -0.13] | -6.666 | <0.001 |
|  | sex = male | 0.68 | 0.14 | [0.41, 0.95] | 4.916 | <0.001 |
|  | age^1 | 19.00 | 9.14 | [1.08, 36.91] | 2.079 | 0.038 |
|  | age^2 | 10.43 | 10.84 | [-10.81, 31.68] | 0.963 | 0.336 |
|  | age^3 | 63.43 | 9.98 | [43.86, 83] | 6.353 | <0.001 |
|  | (group = Urban):date | 0.00 | 0.01 | [-0.02, 0.03] | 0.340 | 0.734 |
|  | (sex = male):age^1 | 19.51 | 11.65 | [-3.33, 42.34] | 1.674 | 0.094 |
|  | (sex = male):age^3 | -61.10 | 11.95 | [-84.52, -37.67] | -5.112 | <0.001 |
|  | (sex = male):age^3 | -26.60 | 11.22 | [-48.59, -4.61] | -2.371 | 0.018 |

**Table 1 (cont.).** Linear models including random effects of sleep start, end, and midpoint times, expressed as hours from 00:00 on the date of start of sleep.
|  | Parameter | Coefficient | SE | 95% CI | t | p |
| --- | --- | --- | --- | --- | --- | --- |
| Sleep<br>offset | (Intercept) | 31.4 (2) | 0.12 | [31.17, 31.64] | 261.494 | <0.001 |
|  | group = Urban | -0.58 | 0.13 | [-0.83, -0.32] | -4.443 | <0.001 |
|  | date (years) | 0.03 | 0.01 | [0.01, 0.05] | 2.424 | 0.015 |
|  | photop. length (h) | -0.17 | 0.01 | [-0.19, -0.14] | -12.444 | <0.001 |
|  | day = weekday | -0.21 | 0.03 | [-0.26, -0.16] | -8.011 | <0.001 |
|  | sex = male | 0.56 | 0.15 | [0.26, 0.86] | 3.689 | <0.001 |
|  | age^1 | -27.53 | 9.64 | [-46.43, -8.63] | -2.855 | 0.004 |
|  | age^2 | 35.05 | 11.19 | [13.12, 56.98] | 3.133 | 0.002 |
|  | age^3 | 60.68 | 10.07 | [40.94, 80.41] | 6.026 | <0.001 |
|  | (group = Urban):date | 0.08 | 0.01 | [0.06, 0.11] | 6.428 | <0.001 |
|  | (sex = male):age^1 | 26.99 | 12.03 | [3.42, 50.57] | 2.245 | 0.025 |
|  | (sex = male):age^2 | -86.19 | 12.28 | [-110.26, -62.13] | -7.021 | <0.001 |
|  | (sex = male):age^3 | -19.82 | 11.35 | [-42.07, 2.44] | -1.745 | 0.081 |
| Sleep<br>midpoint | (Intercept) | 27.23 (3) | 0.11 | [27.01, 27.45] | 242.898 | <0.001 |
|  | group = Urban | -0.48 | 0.12 | [-0.71, -0.25] | -4.12 | <0.001 |
|  | date (years) | 0.08 | 0.01 | [0.06, 0.1] | 8.421 | <0.001 |
|  | photop. length (h) | 0.01 | 0.01 | [-0.02, 0.03] | 0.467 | 0.64 |
|  | day = weekday | -0.2 | 0.02 | [-0.24, -0.15] | -9.014 | <0.001 |
|  | sex = male | 0.62 | 0.14 | [0.33, 0.9] | 4.297 | <0.001 |
|  | age^1 | -5.29 | 8.84 | [-22.62, 12.04] | -0.599 | 0.55 |
|  | age^2 | 28.04 | 10.11 | [8.23, 47.85] | 2.774 | 0.006 |
|  | age^3 | 66.76 | 8.98 | [49.16, 84.35] | 7.437 | <0.001 |
|  | (group = Urban):date | 0.05 | 0.01 | [0.03, 0.07] | 4.21 | <0.001 |
|  | (sex = male):age^1 | 26.13 | 10.83 | [4.91, 47.35] | 2.413 | 0.016 |
|  | (sex = male):age^3 | -80.52 | 11.06 | [-102.2, -58.84] | -7.28 | <0.001 |
|  | (sex = male):age^3 | -24.61 | 10.14 | [-44.49, -4.73] | -2.427 | 0.015 |

Intercept levels: age = 22 y.o., photop. length = 12 h, group = Rural, date = 2018/7/1, sex = female, type of day = free day.
Clock time conversion: (1) 22:57; (2) 7:24; (3) 3:14

**Table 2.**
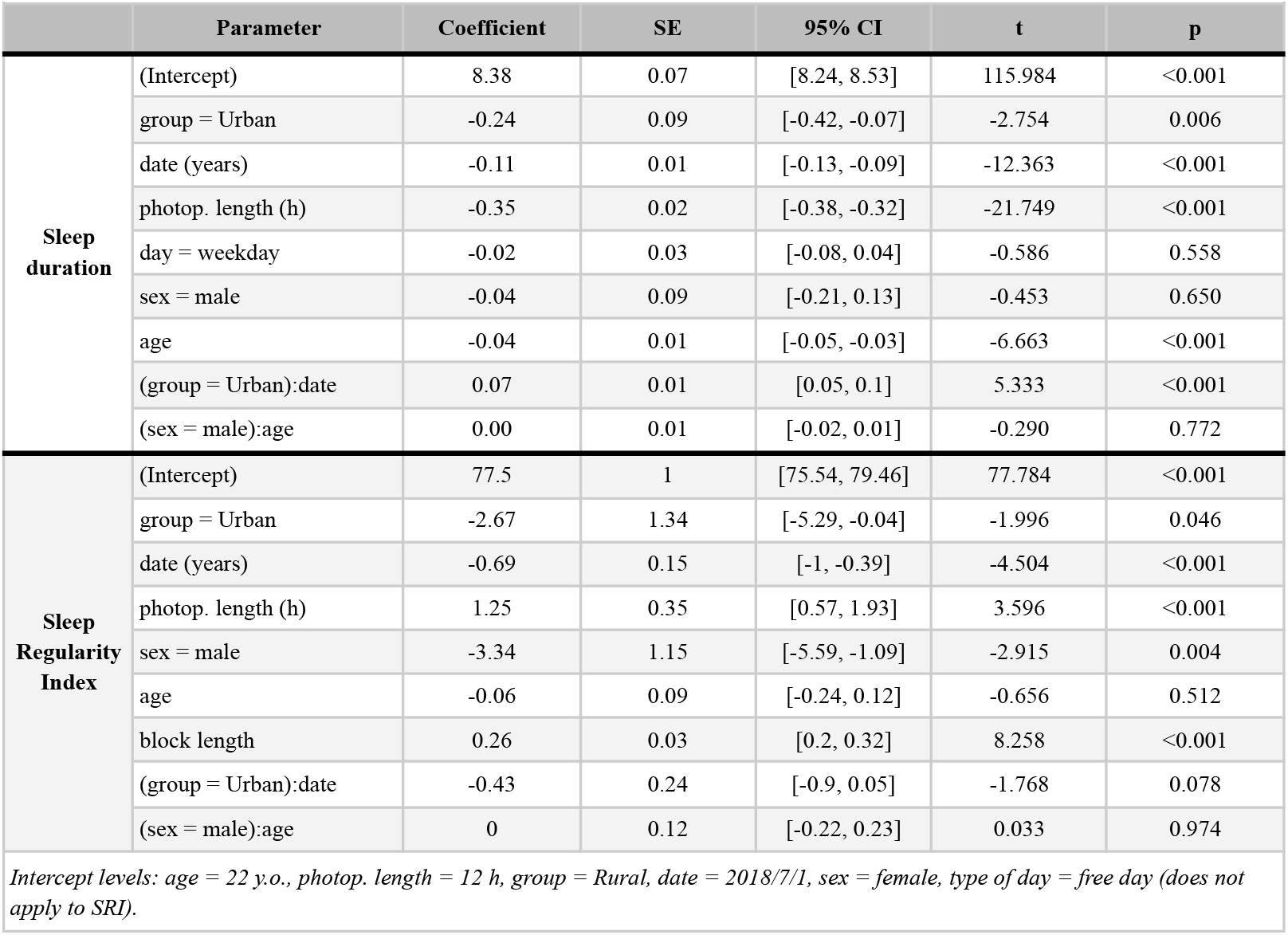
Linear models including random effects of sleep duration (h) and sleep regularity (SRI units)

|  | **Parameter** | **Coefficient** | **SE** | **95% CI** | **t** | **p** |
| --- | --- | --- | --- | --- | --- | --- |
| **Sleep duration** | (Intercept) | 8.38 | 0.07 | [8.24, 8.53] | 115.984 | <0.001 |
|  | group = Urban | -0.24 | 0.09 | [-0.42, -0.07] | -2.754 | 0.006 |
|  | date (years) | -0.11 | 0.01 | [-0.13, -0.09] | -12.363 | <0.001 |
|  | photop. length (h) | -0.35 | 0.02 | [-0.38, -0.32] | -21.749 | <0.001 |
|  | day = weekday | -0.02 | 0.03 | [-0.08, 0.04] | -0.586 | 0.558 |
|  | sex = male | -0.04 | 0.09 | [-0.21, 0.13] | -0.453 | 0.650 |
|  | age | -0.04 | 0.01 | [-0.05, -0.03] | -6.663 | <0.001 |
|  | (group = Urban):date | 0.07 | 0.01 | [0.05, 0.1] | 5.333 | <0.001 |
|  | (sex = male):age | 0.00 | 0.01 | [-0.02, 0.01] | -0.290 | 0.772 |
| **Sleep Regularity Index** | (Intercept) | 77.5 | 1 | [75.54, 79.46] | 77.784 | <0.001 |
|  | group = Urban | -2.67 | 1.34 | [-5.29, -0.04] | -1.996 | 0.046 |
|  | date (years) | -0.69 | 0.15 | [-1, -0.39] | -4.504 | <0.001 |
|  | photop. length (h) | 1.25 | 0.35 | [0.57, 1.93] | 3.596 | <0.001 |
|  | sex = male | -3.34 | 1.15 | [-5.59, -1.09] | -2.915 | 0.004 |
|  | age | -0.06 | 0.09 | [-0.24, 0.12] | -0.656 | 0.512 |
|  | block length | 0.26 | 0.03 | [0.2, 0.32] | 8.258 | <0.001 |
|  | (group = Urban):date | -0.43 | 0.24 | [-0.9, 0.05] | -1.768 | 0.078 |
|  | (sex = male):age | 0 | 0.12 | [-0.22, 0.23] | 0.033 | 0.974 |

*Intercept levels: age = 22 y.o., photop. length = 12 h, group = Rural, date = 2018/7/1, sex = female, type of day = free day (does not apply to SRI).*

Figure 2 and Table S2 present the model-predicted changes in sleep variables for each group under a 12-h photoperiod length for a 22-year-old (the median of ages across all campaigns), while averaging for sex and type of day, for the 2014 to 2024 ten-year span. Models indicate similar delays in the times of start of sleep of 1.36 [1.15-1.58] h (t = 12.611, p < 0.001) for the rural group and of 1.41 [1.17-1.65] h (t = 11.568, p < 0.001) for the urban group. Evolution of sleep end times was asymmetrical between groups: while the rural group changed delayed wake-up times slightly over ten years (0.27 [0.05-0.48] h, t = 2.420, p = 0.016), the urban participants delayed theirs substantially (1.11 [0.88-1.35] h, t = 9.213, p < 0.001). In consequence, changes in sleep duration were also different between communities. While the rural group displayed the loss of over a full hour of sleep (1.1 [0.93-1.28] h, t = 12.336, p < 0.001), this reduction was smaller in the urban group (0.37 [0.16-0.59] h, t = 3.386, p < 0.001). This asymmetry in the evolution of wake-up times also implicated that the urban group showed the larger shift in their midpoint of sleep, with a delay of 1.3 [1.09-1.51] h, as compared to the 0.83 [0.63-1.02] h delay of the rural group. Sleep regularity, measured by the Sleep Regularity Index, showed a significant decrease in both groups across time. Between 2014 and 2024, the model predicts a loss of 6.9 [3.9-10.0] units in the rural population (t = 4.489, p < 0.001) and of 11.2 [7.5-15.0] units in the urban group (t = 5.917, p < 0.001).

**Fig. 2.**
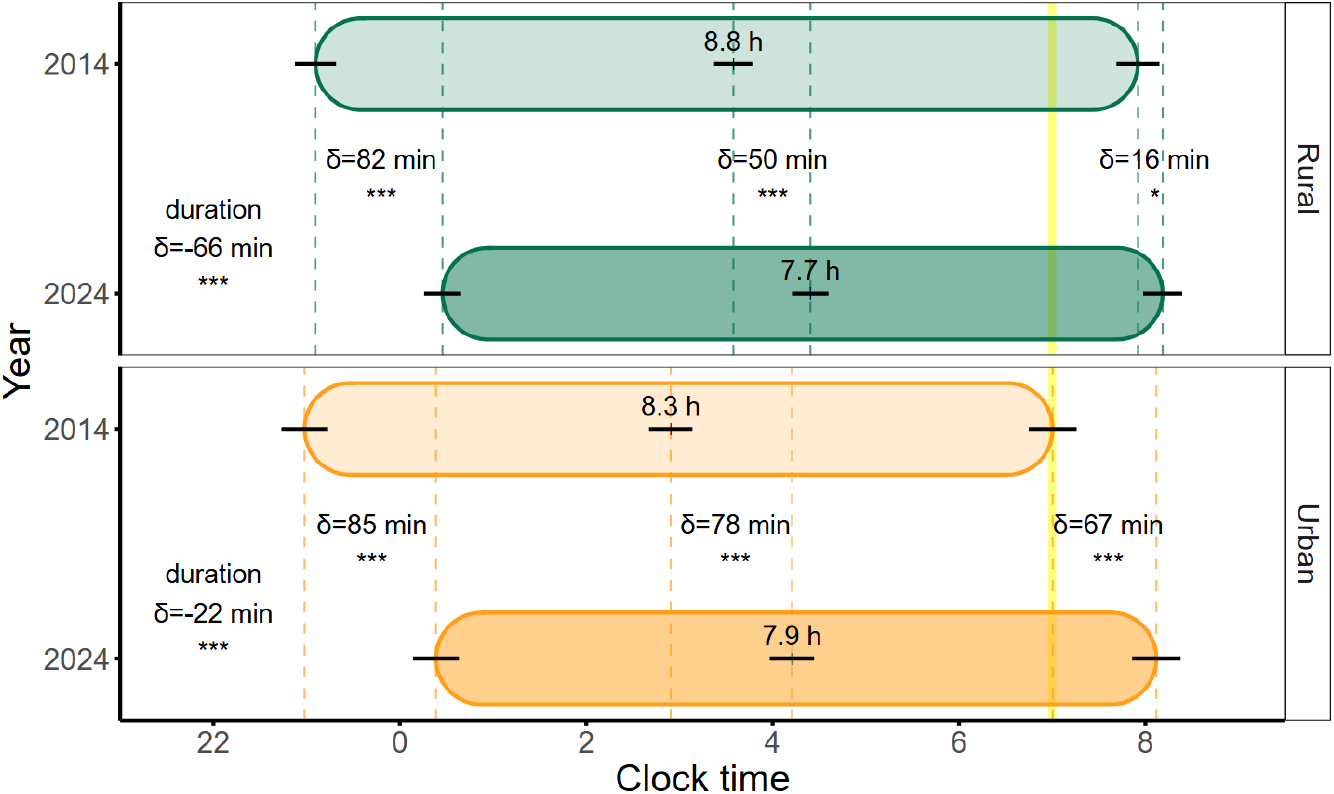
Predictions of sleep patterns for 2014 and 2024 based on linear models including random effects under a 12 h photoperiod (averaged for sex and type of day, and with age set at 22 y.o., the median across all studies). Vertical dashed lines were drawn for the start, end, and midpoint of sleep, and horizontal black lines represent the standard error of these predictions. Predicted sleep duration in hours is also indicated within each box. Predicted differences in minutes are indicated for all sleep variables, and asterisks indicate statistically significant changes across the modelled 2014-2024 span (see Table S2 for the full results of the analyses). The yellow vertical line represents the approximate time of dawn. * p<0.05, *** p<0.001 (Tukey-corrected for multiple comparisons)

The 2012 to 2024 period involved important transformations in these Toba/Qom communities. First, starting around 2016, electricity was progressively introduced in rural communities in the Vaca Perdida and other neighboring areas (Fig. 1.A and Fig. S1). Secondly, while to our knowledge not a single participant from either community under study nor anyone in their families owned a smartphone by 2018, about half of the participants in our studies had their own smartphone, and about the same had access to a TV and/or a computer at their homes by October, 2024. Moreover, even in very remote rural areas, where cell phone coverage is extremely limited, some participants are able to access the internet through free public Wi-Fi hotspots.

We set out to test whether either of these two events could support the observed changes in sleep timing across this timespan, based on their potential for extending activity in the evening hours and thus delaying the start of sleep. To do this, we fitted a new linear model in which every sleep event was labeled as recorded either under conditions of access to electricity or not, and whether it was recorded during the “smartphones and internet era,” an umbrella term for all data from 2023 onwards (Table S3). This model revealed that the “smartphone era” accounted for a delay of 0.40 [0.13-0.67] h in the average time of sleep start (t = 2.913, p = 0.004), while the access to electricity factor did not explain a significant amount of the observed delay in sleep times. Additionally, across the time span there was a delay in sleep start times of 0.09 [0.04-0.14] h per year (t = 3.454, p < 0.001); adding up to about a full hour in total — i.e. the unexplained portion of the ∼1.4 h delay in the previous analysis (see Fig. 2). These results evidence a continuous process of a shift of sleep to later times.

## Discussion

Our analysis reveals the rapid evolution of sleep habits in two Toba/Qom communities during early industrialization phases. Based on 12,000 recorded sleep events across twelve years, our models reveal drastic changes in sleep timing, including a delay in sleep onset of around one and a half hours, and a reduction in daily sleep duration of over an hour in rural communities. Our results mimic —in a timespan an order of magnitude shorter— what was likely a century-long historical change in sleep phase and duration as humans transitioned from preindustrial to industrial and postindustrial environments ^14^. Our findings also suggest there are other factors driving sleep changes beyond the availability of electricity and the use of electric light, and even smartphones, computers, or TV sets. These results underscore the importance of monitoring the impact of the introduction of modern features on the lifestyle of preindustrial communities.

In the time we studied these communities, we detected similar delays in the time of sleep start across groups, but only the urban community displayed a comparable delay in the offset of sleep. Rural communities, who live under even more limited conditions than their counterparts in the city of Ingeniero Juárez, may not be able to extend their sleep much later in the morning either due to being more exposed to natural elements (sunlight, temperature), to the need to start with sustenance activities (e.g. harvesting firewood, managing livestock, gathering), or to other necessities related to their isolated rural environments (e.g. distance to health centers or convenience shops, lack of transportation options). This contrast may explain the more drastic reduction in sleep duration in the rural group in the face of similar shifts in the timing of sleep. That is, the large shift in the timing of sleep start appears to be the most relevant culprit driving changes in overall sleep behavior in these communities.

Internet access and devices such as smartphones, however limited in supply, allow Toba/Qom communities to connect to a world beyond their local environment, for example, through social media and video platforms. The adoption of these technologies in these communities has been very rapid. To the best of our knowledge, smartphones were virtually unavailable by 2018, yet by 2024, half of the Toba/Qom participants owned a smartphone, and about the same had access to a TV or a computer in their homes, reflecting a swift shift toward digital connectivity. A public internet antenna was installed as recently as 2022 in Isla García, providing free internet access to the community; one family in Vaca Perdida even had a satellite internet connection. These technological advances likely favor the delay of sleep and, whenever the time for waking up cannot be further delayed, the reduction of sleeping time by providing almost unlimited entertainment for the night hours. However, our models indicate that this era of the introduction of smartphones and the internet can only explain about 25 minutes of the over 80 minutes of observed delays in the start of sleep. Moreover, this delay in sleep times cannot be driven merely by late-night smartphone users. A recent study on these communities revealed that users of smartphones display only a 12-minute delay of sleep compared to non-users ^15^, making it impossible for just these users to explain the population-level changes reported here. But there is more to this technological adoption than the substitution of sleep for electronic device use or exposure to electric light sources. Electronic devices such as smartphones, computers, and TVs, along with access to the internet, can open a door into the “modern” lifestyle, of which nightlife is a central element. This is particularly relevant in Argentina, where, for instance, the TV “prime time” extends from 21:00 to midnight and football matches can start as late as 22:00 ^16^. This phenomenon merits further research incorporating Toba/Qom self-perceived experiences regarding access to electricity and electric light sources, as well as the roles of the internet, the TV, and other electronic devices.

Sleep loss is considered a modern world epidemic with significant consequences for public health, performance and safety, learning and memory, and overall economic impact ^17–20^. Lack of sleep can affect health at several levels, including, but not limited to, cardiovascular, metabolic, immune, and mental health ^21,22^, and is a strong predictor of all-cause mortality ^23^. Although our models predict an average sleep duration still within the consensus recommended sleep duration range for these ages ^24^, we note that these recommendations are aimed at people in modern developed societies. For a deeply underprivileged population such as the Toba/Qom, which experiences reduced access to health services, dwells in extremely harsh rural environments, and suffers from malnutrition ^25^, such a sharp reduction of sleep time could be more challenging for health and wellbeing than in developed postindustrial settings. Alternatively, the time that the Toba/Qom people lose from sleep for activities such as learning, communal participation, or even leisure (including the use of electronic devices) may offer additional benefits to their quality of life. More research is needed to assess whether this reduction of sleep time in this population is associated with changes in physical health, psychological wellbeing, and overall quality of life.

The observed reduction in sleep regularity in both communities may be more concerning. Sleep regularity has gained attention recently as a marker of sleep health ^26^, and has been proposed as a more precise predictor of the negative impacts of poor sleep than sleep duration itself ^27^. Delayed sleep has already been associated with low sleep regularity in hyper-industrialized societies ^28^, and in a study combining data from adolescents in the Toba/Qom and in rural communities of Southern Brazil, we already reported that sleep regularity was reduced in students exposed to electric light in the evening hours ^12^. Further investigation should determine whether this change is driven by the mere access to electric light and other electric appliances and devices at home, and/or if it is correlated with changes in evening habits that involve the whole community.

Numerous studies of preindustrial communities support the notion that electricity and other postindustrial environmental factors impact sleep timing ^5^. However, the hypothesis that industrialization has necessarily shortened sleep remains under debate. A recent meta-analysis, for instance, suggests that people in industrialized societies sleep longer than those in preindustrial settings, and attributes modern sleep-related health issues not to shorter sleep but to heightened circadian disruption ^3^. However insightful, these observations need to be carefully weighed, considering the inherent diversity of pre-industrial and industrialized groups, without overlooking the unique environmental and sociocultural factors that can shape sleep. Our longitudinal study of the Toba/Qom communities provides a rare opportunity to observe the real-time impact of modernization, including the acquisition of electricity and related technologies. Our data reveals that the modernization of Toba/Qom communities correlates with a drastic reduction in sleep duration and an increased circadian disruption —marked by a reduced sleep regularity and the further decoupling of sleep onset from sunset. We should also note that the transition from harsh preindustrial conditions to an electrically lit postindustrial environment can also have beneficial effects on sleep, for example, by improving comfort and sense of safety at bedtime ^29^. This underscores the intricate ways in which the environment and modernization interact to shape human sleep patterns ^30^.

Some limitations in our study need to be addressed. First, it should be noted that the recordings in our dataset have been obtained using different devices and algorithms. Because of the long timespan of our study, this was an obstacle hard to overcome. Of note, the Actiwatch devices used from 2012 to 2018 do not provide raw accelerometric data, as the Axivity AX3 devices do, precluding us from using a common algorithm to extract sleep timing. However, we are confident that the different methods are sufficiently comparable as they are all based on longitudinal wrist-accelerometry and the sleep estimation was performed following well-tested protocols, based on similar criteria of activity/inactivity and aided by sleep logs to detect unreliable estimations. Moreover, our tests indicate that sleep estimation from AX3 and Actiwatch Spectrum devices are comparable (see Fig. S2), especially considering the large amount of data available. Given the magnitude of the effects estimated by our statistical models, it is improbable that slight differences in the measuring tool would artificially generate the changes we observed.

Second, thanks to the improvement of fieldwork conditions, in terms of the available equipment and our relationship with the communities, our dataset is much denser in recent years, compared to the first four years of work. While this may pose a limitation in the model’s precision for those early years, the use of linear models, including random effects accounting for sex, age, and repeated measures, is robust enough to assess the dynamics of sleep across the studied period in these communities. Importantly, the lower number of participants in the rural community before the appearance of electricity may have contributed to the lack of a statistically significant effect of electricity on sleep onset in our analysis. Obviously, this result does not mean that the appearance of electricity was not critical for the sleep delay and shortening we observed; electricity not only allows for artificially lit evenings but also for internet hotspots, cell phones, TVs and other sleep disruptors.

Our work highlights the challenges and multifaceted impacts of technological innovation on human behavior and health, especially among underprivileged populations. While these communities have finally received the innumerable benefits of access to electricity, smartphones, and the internet, they remain significantly behind in accessing many of the advantages of modern, developed societies. This disparity may exacerbate the negative consequences of delayed and shortened sleep in these communities, which seem to be inescapable features of modern lifestyle. In this sense, the Toba/Qom case we present exemplifies what has been described as *“cultural arrhythmia,”* i.e., an imbalance between the pace of changes in human lives and their surrounding habitats ^31^. In this sense, it is imperative that such transformative processes consider the unique features and conditions of each community, ensuring they enhance health and overall well-being while proactively addressing any potential challenges. Our results highlight the urgent need to design specific public health policies and awareness campaigns to protect sleep health in groups that might be particularly vulnerable to these changes.

## Methods

### Participants and ethics

All the activities described in this work, past and recent, were reviewed and approved by the Internal Review Board from the University of Washington’s Human Subject Division and followed the Declaration of Helsinki. The research did not involve any risk to participants or researchers.

The participants belonged to three Toba/Qom communities in Formosa, a northern province in Argentina, all of which share a recent historical past and are closely related culturally and ethnically ^8^:

1. *Barrio Toba*: a community located adjacent to Ingeniero Juárez (23°47′ S, 61°48′ W), a town with close to 20,000 inhabitants, including the Toba/Qom people. This community has had 24/7 access to electricity at their homes throughout the time we surveyed them. People in this community settled adjacent to Ingeniero Juárez after migrating from the northern Toba/Qom communities in the 1990s.
2. *Vaca Perdida*: a rural community of a few hundred members located ∼60 km north of Ingeniero Juárez (23°29′ S, 61°38′ W). This community slowly gained access to electricity when the local government started introducing it in 2016 and has since had regular access to electricity and, more recently, paid internet access.
3. *Isla García*: this small rural community (50-70 people) is named after the family that founded it. Since we started working with them, the García family and their relatives have moved through three different locations in the vicinity of Vaca Perdida. The first year they were able to consistently access electricity was 2017, and since 2018, they have remained at their current location with full access to electricity. In 2023, the local government installed a public internet antenna on the community grounds.

Consent was obtained orally from every participant after detailing the study procedures in Spanish, with the help of a Toba/Qom translator whenever necessary. For subjects under 18 years of age, consent as well as assent by one of the parents were obtained. Participants were not monetarily compensated for their participation in any of these studies. In appreciation of their participation, they received different gifts, usually clothing and daily-use items.

It is important to note that these communities engage in very similar social behavior and cultural activities. Children and adolescents attend school during daylight hours, although the time for each age group may differ slightly between communities. Schools are typically located within walking distance of the houses. Most adults are formally unemployed and rely economically on occasional gigs and government assistance. Of note, substance abuse disorders are rare among these communities, in part due to their Anglican religious practices; indeed, selling alcohol is forbidden in Vaca Perdida. These groups are among the most underprivileged in Argentina. Recently, the National Institute of Statistics and Census (INDEC) classified communities in this area as “very low income/poor.” Socioeconomic differences between families are negligible and thus were not included as a covariate in analyses.

Classifying rural and urban communities includes considering factors such as access to electricity, built housing, and market and labor integration. Beyond their history of access to electricity, rural communities are far from health, continued education, and administrative centers, grocery shops, and banks, and usually lack transportation mediums of their own to travel the challenging ∼60 km road necessary to access these in Ingeniero Juárez. The “rural” group in this paper is represented by the aggregated data of both Vaca Perdida and Isla García participants. The Toba/Qom community in Barrio Toba will be assigned the “urban” label in this paper, due to being settled within an urbanized area, even if their access to these urban amenities may be limited due to their economic situation.

#### Survey of electronic devices access

Starting in the 2023 Spring campaign, when signing up, each participant was asked whether they owned a smartphone. We followed up by asking participants if they had access to a TV, a computer or tablet device, and/or an internet connection at their homes.

### Data acquisition, treatment, and analysis

Data was collected at different points between 2012 and 2014, 2016 and 2018, and 2023 and 2024, and comprises a total of 12,307 recorded sleep events from 156 participants. Data was recorded mostly in the late fall or early winter, and in the late spring or early summer, to better capture seasonal differences. Table S1 details the number of participants, their age and sex composition, and the number of recorded events according to community, year, and season of collection. One-hundred and fifty-six (156) unique subjects participated in the study (94 from the rural group, 63 from the urban group; one participant moved from the urban group to the rural group after 2018; 52% females). Table S4 indicates the total number of participants in each field campaign, split by study group, along with the number of participants recorded for the first time in each. Table S5 shows the distribution of records across participants, in terms of the number of campaigns in which they participated; around 50% of the participants were recorded at two or more timepoints. Fig. 1.B further illustrates the longitudinal nature of our study.

All the data manipulation and analysis were carried out in the *R* language ^32^ (version 4.5.1) on the RStudio platform ^33^ unless specifically indicated.

#### Sunlight and light pollution data

Solar data for the Ingeniero Juárez and Vaca Perdida geographical regions was obtained using the *suncalc* package ^34^. For every date present in the dataset, we collected the sunrise and sunset times. These were used to calculate the natural photoperiod for each date, as well as for the filtering of nightly sleep episodes that are described below.

Longitudinal values of night light pollution were obtained by the Day/Night Band of NASA’s Black Marble Group’s Visible Infrared Imaging Radiometer Suite instrument (VIIRS DNB, zero point corrected). Square one km^2^ areas surrounding the location of the studied communities were selected using the Radiance Light Trends application (https://lighttrends.lightpollutionmap.info/), and values were downloaded for all months from April 2012 to July 2024. Satellite images of the regions of interest superimposed with VIIRS radiance data through the years were obtained from Jurij Stare’s Light Pollution Map (www.lightpollutionmap.info).

#### Sleep estimation

Sleep was assessed by wrist actigraphy, and participants also completed daily sleep logs throughout each recording session. Participants wore the actigraphs on their non-dominant wrists, and locomotor activity data was always collected in one-minute bins. Individual recordings were visually examined before proceeding to the sleep estimation to confirm data integrity and discard low-quality recordings (e.g. fragmented data, frequent missing periods of data, frequent saturated counts of activity).

Three different wrist-actigraphic devices were used across the years, and consequently, the methods used for sleep detection also differed:

1. Participants in the studies before 2014 wore the Actiwatch-L device from Mini Mitter (Bend, OR); data was analyzed for sleep detection, aided by sleep logs, using the Sleep Analysis package of the Mini Mitter Software (version 3.4), as described in ^9^.
2. Participants recorded between 2014 and 2018 were equipped with the Actiwatch Spectrum Plus loggers from Respironics (Bend, OR); sleep detection was performed using the Philips Respironics Actiware software V.6.0.9, as described in ^35^.
3. The newly reported data in this paper, collected since 2023, were obtained with the AX3 devices from Axivity (Newcastle upon Tyne, UK), which are equipped with a tri-axial accelerometer. Devices were set at 50hz with a range of 8g. Sleep detection was performed using the GGIR software for R ^36^ using the implemented Van Hees algorithm ^37^. A minimum of 16 hours of wear-time was required for each day of record.

Algorithm-based sleep detection from data cohorts 2 and 3 above was contrasted with individual sleep logs non-exhaustively to assess the overall rationality of estimations.

We are aware that the use of different devices and algorithms imposes an important limitation on the precision of our analysis. However, we have strong reasons to trust our overall conclusions: firstly, daily sleep logs were used to check the reliability of the detected sleep bouts, and secondly, all three combinations of devices and algorithms estimate sleep based on criteria of sustained activity/inactivity on one-minute bins of actigraphic recordings.

Moreover, we performed a comparison of the sleep/wake estimations of the Actiwatch Spectrum Plus and the AX3 devices by comparing the data obtained from 22 non-Toba/Qom volunteers using both devices on their non-dominant wrist concomitantly for 7 to 14 days. We performed a Bland-Altman analysis ^38^ based on 206 hundred sleep events, selected with the same criteria used for the Toba/Qom data (described in the next subsection). Due to the non-normality of the differences, we performed a non-parametric analysis of the results. The analysis revealed that differences between the AX3/GGIR and Actiwatch/Actiware methods can take a wide range of values, but that the median of these differences is very small for both sleep start and end time, as well as for duration. Differences between methods, expressed as AX3/GGIR minus Actiwatch/Actiware: for sleep start, mean = 16.5 min, median = 1.4 min; for sleep end, mean = -0.9 min, median = 2.6 min; for sleep duration, mean = -17.5 min, median = 1.2 min. See Fig. S2 for more details. All of these suggest that these two methods should provide very similar estimates for a large enough sample such as ours, comprising over ten thousand data points. Moreover, the observed mean differences between methods are much smaller than the observed effect sizes.

#### Sleep data treatment and removal of outlier data

The whole 2012-2024 dataset of sleep events was filtered first to keep only nightly sleep events. Every sleep event starting outside the hours from dusk till dawn on any date was removed from the dataset. This step removed only 181 sleep events (1.5% of the total sleep events). While actigraphy performs poorly for the detection of naps, daytime sleep is rare in these Toba/Qom communities. According to the analysis of sleep logs, the proportion of Toba/Qom that reported naps decreased from 75% in 2012 to 36% in 2024 (Fig. S3.A). The frequency of reported siestas was very low and changed from a median of 2.3 naps per week in 2012 to only one nap per week in 2024 (Fig. S3.B).

We then proceeded to detect events that could be considered outliers based on their duration (i.e. too long or too short events). This approach allowed us to remove data points that could represent either artifacts of the sleep detection method, or sleep events from “non-representative” nights (e.g., during travel or while sick). The outlier analysis was performed using the median absolute deviation (MAD) method ^39^ from the *Routliers* package ^40^, using a threshold of 2.5 MADs (lower limit = 3.49 h, upper limit = 12.44 h). This step removed an additional 506 (4.1%) sleep events from the dataset. The final dataset on which the analysis was performed had a size of 11,620 sleep records.

#### Sleep regularity

Sleep regularity was evaluated using the Sleep Regularity Index (SRI). The SRI, proposed by Phillips et al. 2017 ^28^, represents a re-scaled measure of the probability that a participant is at an equal state of wake or sleep at two time points separated by 24 hours on any series of at least seven consecutive night sleep events. A person who always wakes up and falls asleep at the same time every day will score 100, while a person who randomly falls asleep and wakes up at any time of the day will score 0.

First, we detected blocks of consecutive nightly sleep events for each participant. Subjects were considered to be asleep within the recorded sleep bout and awake outside of it. Starting at the time of the first recorded sleep start and in one-minute bins, we calculated the SRI for each individual block. Only scores for sets of at least seven consecutive recordings were included in the analysis. It is important to note that for the SRI analysis, the filter step of outliers based on sleep duration described above was purposefully not performed, as we preferred to prevent the loss of the continuity of recordings, even if that meant including potentially inaccurate records of sleep.

#### Statistical analysis

To evaluate the evolution of night sleep timing in these communities, we fit linear models including mixed-effects using the *lme4* package ^41^. Fixed factors included a binary rural/urban factor, a continuous date variable, photoperiod length, type of day (before a workday or before the weekend or a holiday; excluded in the analysis of the SRI), sex, and age. Interaction terms for group and date, as well as sex and age, were also included. Exclusively for the analysis of the sleep start, midpoint, and end variables, the age factor was included as a 3-grade polynomial term to account for the known non-linear progression of sleep timing across adolescence and adulthood ^42^. Individual identities were included as a random factor to account for repeated measures. Statistical significance was tested through a Type III ANOVA as implemented in the *parameters* package ^43^. Below is an example of a *lme4* model equation in this study, illustrating the interaction terms and general syntax:

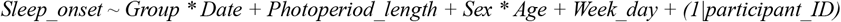

Intercept levels for all linear models were set as follows: age = *22 y*.*o*. (the median age across all participants), photoperiod length = *12 h*, group = *Rural*, date = *July 1st 2018 (i*.*e. 2018*.*5)*, sex = *female*, type of day = *free day*.

The assumptions of the models were evaluated using the *check_model* function from the *performance* package ^44^. We tested the following features: 1) *posterior predictive check* (are real values far from modelled values?), 2) *linearity*, 3) *homogeneity of variance*, 4) *normality of residuals*, 5) *normality of random effects*, and 6) *influential observations* (are there too many outliers?).

To evaluate post-hoc contrasts between years or groups, we used the *emmeans* package ^45^. The tools in this package return the values of the dependent variable predicted by the models based on given independent variables, and test contrasts between levels (p-values are Tukey-corrected for multiple comparisons). A ten-year span between 2014 and 2024 was used when comparing between years to allow for an easier appreciation of the magnitude of the results. We decided to set the modelled age at 22 years old, which is the median age of participants in the dataset across all recording campaigns. It is important to note that age was not included as an interacting factor with groups, years, or photoperiod length in our models, and thus while the absolute predicted values can change according to their dynamics through age, the differences in the contrasts between the aforementioned factors do not change regardless of the set age value. In other words, had we applied our model to a 40-year-old participant, the delays or advances in sleep timing between the two communities would be the same, even when absolute sleep times may be different.

## Acknowledgments

The authors extend their gratitude to the participants from the Toba/Qom communities of Ingeniero Juárez, Vaca Perdida, and Isla García. Their welcoming generosity has been invaluable to our research team. We would like to thank especially community leaders whose assistance on-site has been essential for our work. Additionally, we are grateful to Hotel Indiano in Ingeniero Juárez for their hospitality and support.

We are thankful to Professors Claudia Valeggia and Eduardo Fernández-Duque (Yale University), who introduced our group and were key to establishing our ties to these communities. Marcelo Rotundo and Fundación ECO (Formosa) provided invaluable help during field campaigns throughout the years.

Finally, the authors thank all the researchers who participated in fieldwork across all these years.

This work has been supported by funding from these agencies and institutions:

- National Institutes of Health, Grant # R01HL162311
- Universidad de San Andrés, Program of Support to Research

## Author contributions

Data collection: LPC, IS, LLT, MFC, CRGP, VYZ, HOD

Data analysis: LPC, IS, MDM, DAG, HOD

Writing—original draft: LPC

Writing—review & editing: all authors

Funding acquisition: LPC, DAG, HOD

## Data and materials availability

sleep data will be made available in a public database; a link will be provided upon acceptance of the manuscript.

## SUPPLEMENTARY MATERIAL

Contains three (3) supplementary figures and five (5) supplementary tables.

**Figure S1.**
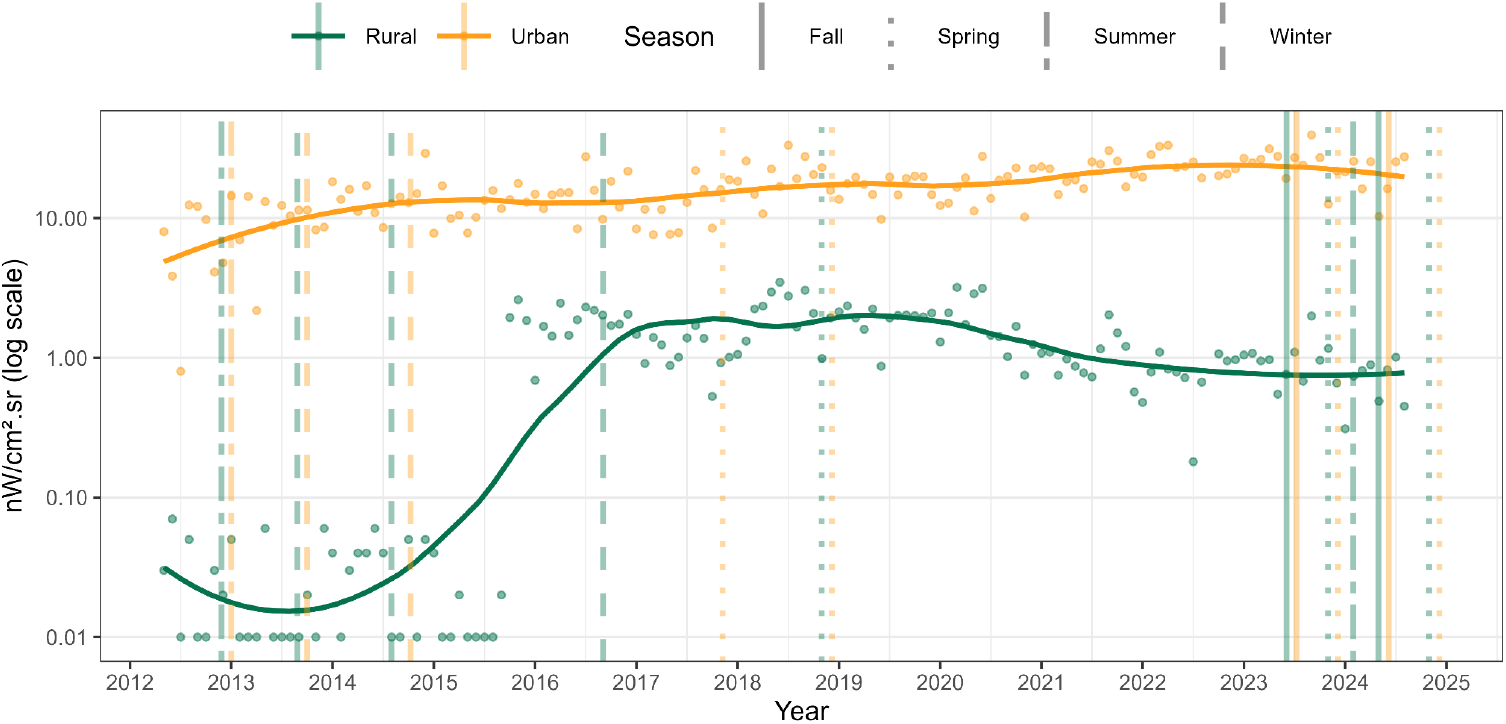
Evolution of night light pollution as a proxy of the introduction of electricity in the study areas. Monthly night radiance DNB levels measured by NASA’s VIIRS instrument for 4 km^2^areas around Barrio Toba in Ingeniero Juárez (urban) and the center of Vaca Perdida (rural). The sharp increase in 2016 marks the beginning of work to install the electricity grid in the rural area. Vertical lines indicate approximate times of data collection since 2012; lines were purposefully spaced during field campaigns that included both groups for easier visualization.

**Figure S2.**
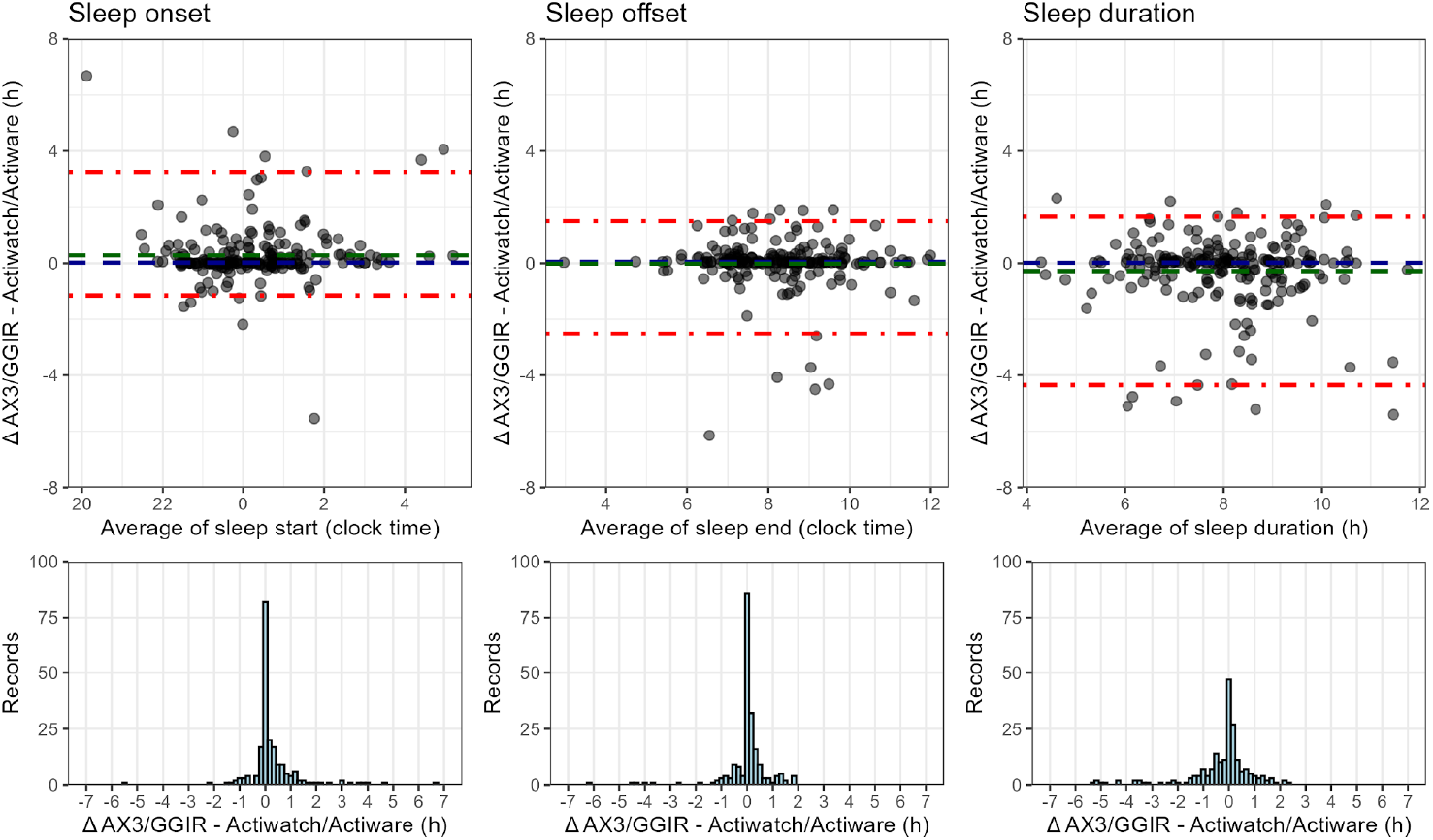
Comparison of the Axivity AX3+GGIR and Actiware Spectrum Plus+Actiware methods for the detection of sleep from wrist-actigraphy recordings. Data consists of 206 detected sleep events (subject to the same cleaning procedure as the data from the main study) from 22 subjects wearing both devices on their non-dominant wrist from 7 to 14 days. Top: Bland-Altman analysis of the differences. The red dashed lines indicate the margin containing 95% of the observations; the blue dashed lines represent the mean of the differences, while the blue dashed line represents the median. Bottom: Distribution of the observed differences.

**Figure S3.**
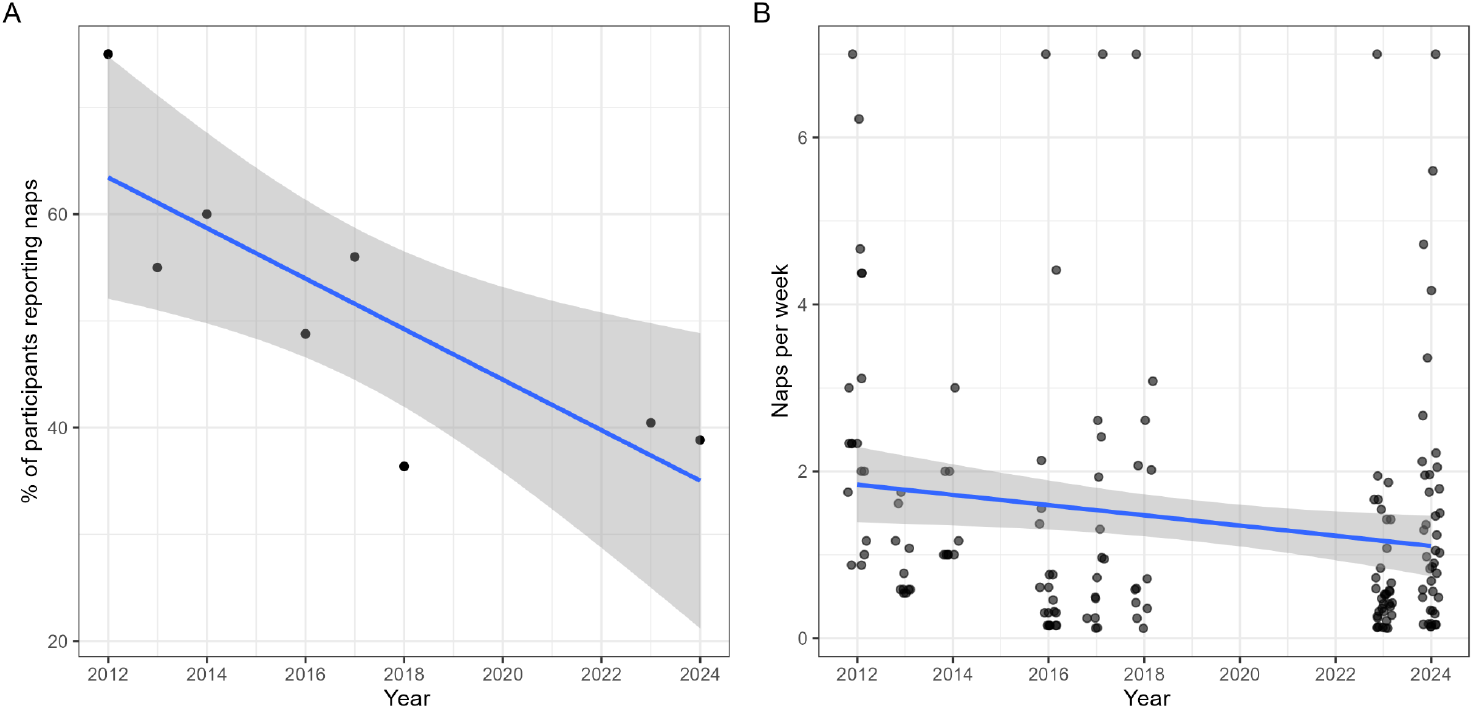
Evolution of naps in the Toba/Qom communities studied. **(A)** Percentage of participants that reported at least one daytime nap during a recording period, split by year. **(B)** Frequency of daytime naps reported (in naps per week), by year. Lines and shades represent linear regressions of the datapoint, with added standard errors.

**Table S1.**
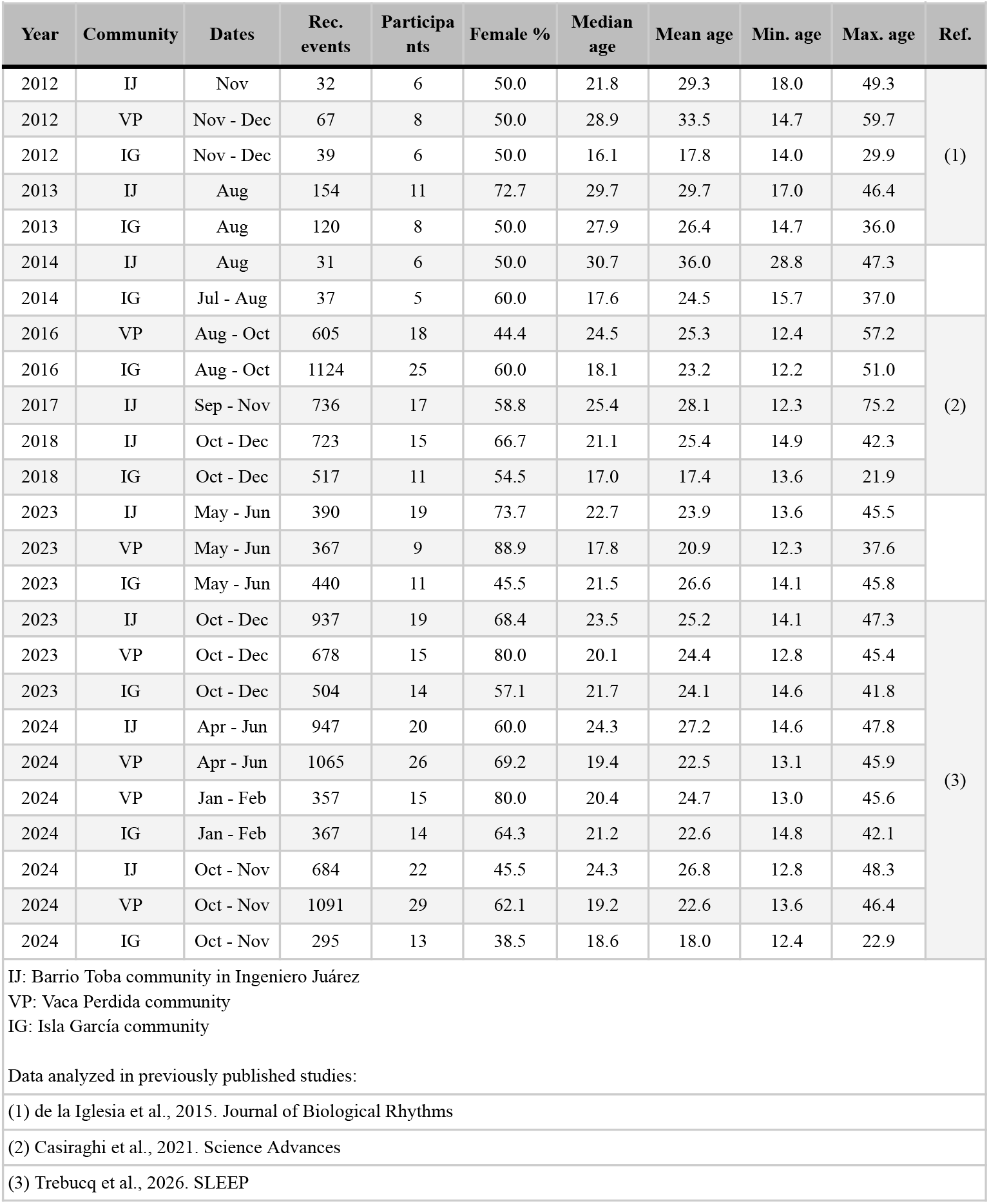
Demographic characteristics for all participants in the dataset.

**Table S2.** Linear model predictions for years 2014 and 2024 for each community, for a 22 y.o. participant under a 12 h photoperiod, averaged for sex and type of day (not applicable to SRI).

| Variable | Group | Year | Predicted | 95% CI lower | 95% CI upper | Diff. | 95% CI lower | 95% CI upper | t ratio | p value |
| --- | --- | --- | --- | --- | --- | --- | --- | --- | --- | --- |
| Onset (clock time) | Rural | 2014 | 23.09 | 22.87 | 23.32 | -1.36 | -1.58 | -1.15 | -12.611 | <0.001 |
|  |  | 2024 | 0.46 | 0.26 | 0.65 |  |  |  |  |  |
|  | Urban | 2014 | 22.98 | 22.73 | 23.23 | -1.41 | -1.65 | -1.17 | -11.568 | <0.001 |
|  |  | 2024 | 0.39 | 0.14 | 0.63 |  |  |  |  |  |
| Offset (clock time) | Rural | 2014 | 7.92 | 7.69 | 8.15 | -0.27 | -0.48 | -0.05 | -2.42 | 0.016 |
|  |  | 2024 | 8.19 | 7.98 | 8.39 |  |  |  |  |  |
|  | Urban | 2014 | 7.01 | 6.75 | 7.26 | -1.11 | -1.35 | -0.88 | -9.213 | <0.001 |
|  |  | 2024 | 8.12 | 7.86 | 8.38 |  |  |  |  |  |
| Midpoint (clock time) | Rural | 2014 | 3.58 | 3.37 | 3.79 | -0.83 | -1.02 | -0.63 | -8.41 | <0.001 |
|  |  | 2024 | 4.41 | 4.21 | 4.6 |  |  |  |  |  |
|  | Urban | 2014 | 2.91 | 2.67 | 3.14 | -1.3 | -1.51 | -1.09 | -12.176 | <0.001 |
|  |  | 2024 | 4.21 | 3.97 | 4.45 |  |  |  |  |  |
| Duration (hours) | Rural | 2014 | 8.8 | 8.64 | 8.95 | 1.1 | 0.93 | 1.28 | 12.336 | <0.001 |
|  |  | 2024 | 7.69 | 7.57 | 7.82 |  |  |  |  |  |
|  | Urban | 2014 | 8.26 | 8.07 | 8.45 | 0.37 | 0.16 | 0.59 | 3.386 | <0.001 |
|  |  | 2024 | 7.89 | 7.73 | 8.05 |  |  |  |  |  |
| Sleep Regularity Index (units) | Rural | 2014 | 78.66 | 76.1 | 81.21 | 6.94 | 3.9 | 9.99 | 4.489 | <0.001 |
|  |  | 2024 | 71.71 | 70.02 | 73.4 |  |  |  |  |  |
|  | Urban | 2014 | 77.7 | 74.61 | 80.8 | 11.22 | 7.49 | 14.95 | 5.917 | <0.001 |
|  |  | 2024 | 66.48 | 64.29 | 68.67 |  |  |  |  |  |

**Table S3.** Linear model of sleep start controlling for the access to electricity and the “smartphones and internet era,” expressed as hours from 00:00 on the date of start of sleep.

| Parameter | Coefficient | SE | 95% CI | t | p |
| --- | --- | --- | --- | --- | --- |
| (Intercept) | 22.81 (1) | 0.13 | [22.55, 23.06] | 176.69 | <0.001 |
| electricity = TRUE | 0.02 | 0.11 | [-0.2, 0.24] | 0.196 | 0.845 |
| smart. era = TRUE | 0.40 | 0.14 | [0.13, 0.67] | 2.913 | 0.004 |
| group = Urban | -0.06 | 0.14 | [-0.34, 0.22] | -0.417 | 0.676 |
| date (years) | 0.09 | 0.03 | [0.04, 0.14] | 3.454 | 0.001 |
| photop. length (h) | 0.19 | 0.02 | [0.16, 0.22] | 12.551 | <0.001 |
| day = weekday | -0.18 | 0.03 | [-0.24, -0.13] | -6.593 | <0.001 |
| sex = male | 0.69 | 0.14 | [0.42, 0.95] | 4.999 | <0.001 |
| age^1 | 15.09 | 9.21 | [-2.96, 33.14] | 1.638 | 0.101 |
| age^2 | 10.02 | 10.81 | [-11.18, 31.22] | 0.927 | 0.354 |
| age^3 | 61.75 | 9.97 | [42.2, 81.29] | 6.193 | <0.001 |
| (group = Urban):date | 0.00 | 0.02 | [-0.04, 0.03] | -0.069 | 0.945 |
| (sex = male):age^1 | 21.80 | 11.64 | [-1.02, 44.63] | 1.873 | 0.061 |
| (sex = male):age^3 | -59.95 | 11.93 | [-83.34, -36.56] | -5.024 | <0.001 |
| (sex = male):age^3 | -24.79 | 11.20 | [-46.74, -2.84] | -2.213 | 0.027 |

*Intercept levels: age = 22 y.o., photoperiod length = 12 h, group = Rural, date = 2018/7/1, sex = female, type of day = free day.*
*Clock time conversion: (1) 22:49*

**Table S4.** Number of participants per field campaign and group, including detail of how many participants were included for the first time in the study in each.

| Campaign | Group | Participants | First time |
| --- | --- | --- | --- |
| 2012 Spring | Rural | 14 | 14 |
| 2012 Spring | Urban | 6 | 6 |
| 2013 Fall/Winter | Rural | 8 | 6 |
| 2013 Fall/Winter | Urban | 11 | 10 |
| 2014 Fall/Winter | Rural | 5 | 1 |
| 2014 Fall/Winter | Urban | 6 | 4 |
| 2016 Winter | Rural | 43 | 32 |
| 2017<br>Winter/Spring | Urban | 17 | 12 |
| 2018 Spring | Rural | 11 | 2 |
| 2018 Spring | Urban | 15 | 8 |
| 2023 Fall/Winter | Rural | 20 | 12 |
| 2023 Fall/Winter | Urban | 19 | 11 |
| 2023 Spring | Rural | 29 | 6 |
| 2023 Spring | Urban | 19 | 4 |
| 2024 Fall/Winter | Rural | 26 | 11 |
| 2024 Fall/Winter | Urban | 20 | 3 |
| 2024 Spring | Rural | 42 | 10 |
| 2024 Spring | Urban | 22 | 5 |
| 2024 Summer | Rural | 29 | 0 |

**Table S5.** Number of participants per field campaign and group, including detail of how many participants were included for the first time in the study in each.

| Time points | Group | Participants | % within group | Total | % from total |
| --- | --- | --- | --- | --- | --- |
| 1 | Rural | 43 | 45.7 | 76 | 48.7 |
|  | Urban | 33 | 52.4 |  |  |
| 2 | Rural | 20 | 21.3 | 32 | 20.5 |
|  | Urban | 12 | 19.0 |  |  |
| 3 | Rural | 4 | 4.3 | 10 | 6.4 |
|  | Urban | 6 | 9.5 |  |  |
| 4 | Rural | 8 | 8.5 | 14 | 9.0 |
|  | Urban | 6 | 9.5 |  |  |
| 5 | Rural | 16 | 17.0 | 18 | 11.5 |
|  | Urban | 2 | 3.2 |  |  |
| 6 | Rural | 1 | 1.1 | 3 | 1.9 |
|  | Urban | 2 | 3.2 |  |  |
| 7 | Rural | 2 | 2.1 | 4 | 2.6 |
|  | Urban | 2 | 3.2 |  |  |

## References

1. Inc, G. (2024). Americans Sleeping Less, More Stressed. Gallup.com. https://news.gallup.com/poll/642704/americans-sleeping-less-stressed.aspx.

2. Matricciani, L., Olds, T., and Petkov, J. (2012). In search of lost sleep: Secular trends in the sleep time of school-aged children and adolescents. Sleep Med. Rev. 16, 203–211. 10.1016/j.smrv.2011.03.005.

3. Samson, D.R., and McKinnon, L. (2025). Are humans facing a sleep epidemic or enlightenment? Large-scale, industrial societies exhibit long, efficient sleep yet weak circadian function. Proc. R. Soc. B Biol. Sci. 292, 20242319. 10.1098/rspb.2024.2319.

4. Bulbulshoev, U., Kassam, K.-A., and Ruelle, M. (2011). Ecology of Time: Calendar of the Human Body in the Pamir Mountains. J. Persianate Stud. 4, 146–170. 10.1163/187471611X600369.

5. Casiraghi, L.P., and de la Iglesia, H.O. (2022). Sleep Under Preindustrial Conditions: What We Can Learn from It. In Circadian Regulation Methods in Molecular Biology., G. Solanas and P.-S. Welz, eds. (Springer US), pp. 1–14. 10.1007/978-1-0716-2249-0_1.

6. Roenneberg, T., Kumar, C.J., and Merrow, M. (2007). The human circadian clock entrains to sun time. Curr. Biol. 17, R44–R45. 10.1016/j.cub.2006.12.011.

7. Yetish, G., and McGregor, R. (2019). Hunter-Gatherer Sleep and Novel Human Sleep Adaptations. In Handbook of Behavioral Neuroscience (Elsevier), pp. 317–331. 10.1016/B978-0-12-813743-7.00021-9.

8. Miller, E.S. (2001). Peoples of the Gran Chaco (Greenwood Publishing Group).

9. de la Iglesia, H.O., Fernández-Duque, E., Golombek, D.A., Lanza, N., Duffy, J.F., Czeisler, C.A., and Valeggia, C.R. (2015). Access to Electric Light Is Associated with Shorter Sleep Duration in a Traditionally Hunter-Gatherer Community. J. Biol. Rhythms 30, 342–350. 10.1177/0748730415590702.

10. Beale, A.D., Pedrazzoli, M., Gonçalves, B. da S.B., Beijamini, F., Duarte, N.E., Egan, K.J., Knutson, K.L., Schantz, M. von, and Roden, L.C. (2017). Comparison between an African town and a neighbouring village shows delayed, but not decreased, sleep during the early stages of urbanisation. Sci. Rep. 7, 5697. 10.1038/s41598-017-05712-3.

11. Moreno, C.R.C., Vasconcelos, S., Marqueze, E.C., Lowden, A., Middleton, B., Fischer, F.M., Louzada, F.M., and Skene, D.J. (2015). Sleep patterns in Amazon rubber tappers with and without electric light at home. Sci. Rep. 5, 14074. 10.1038/srep14074.

12. Peixoto, C.A.T., da Silva, A.G.T., Carskadon, M.A., and Louzada, F.M. (2009). Adolescents Living in Homes Without Electric Lighting Have Earlier Sleep Times. Behav. Sleep. Med. 7, 73–80. 10.1080/15402000902762311.

13. Pilz, L.K., Levandovski, R., Oliveira, M.A.B., Hidalgo, M.P., and Roenneberg, T. (2018). Sleep and light exposure across different levels of urbanisation in Brazilian communities. Sci. Rep. 8, 11389. 10.1038/s41598-018-29494-4.

14. Ekirch, A.R. (2001). Sleep We Have Lost: Pre-industrial Slumber in the British Isles. Am. Hist. Rev. 106, 343–386. 10.1086/ahr/106.2.343.

15. Trebucq, L.L., Moyano, M.D., Coldeira, M.F., Golombek, D.A., De La Iglesia, H.O., Spiousas, I., and Casiraghi, L.P. (2026). Smartphone use is associated with later and shorter actigraphic sleep, and circadian clock delays in an indigenous Toba/Qom community. SLEEP, zsag039. 10.1093/sleep/zsag039.

16. Tassino, B., and Leone, M.J. (2025). Night owls of Rio de la Plata region: Real-life scenarios to understand the biological clock. Neuroscience, S0306452225001368. 10.1016/j.neuroscience.2025.02.022.

17. Anauati, M.V., Gómez Seeber, M., Campanario, S., Sosa Escudero, W., and Golombek, D.A. (2024). The economic costs and consequences of (insufficient) sleep: a case study from Latin America. Eur. J. Health Econ. 10.1007/s10198-024-01733-8.

18. Golombek, D.A., Eyre, H., Spiousas, I., Casiraghi, L.P., Hartikainen, K.M., Partonen, T., Pyykkö, M., Reynolds, C.F., Hynes, W.M., Bassetti, C.L.A., et al. (2025). Sleep Capital: Linking Brain Health to Wellbeing and Economic Productivity Across the Lifespan. Am. J. Geriatr. Psychiatry Off. J. Am. Assoc. Geriatr. Psychiatry 33, 92–106. 10.1016/j.jagp.2024.07.011.

19. Hafner, M., Stepanek, M., Taylor, J., Troxel, W.M., and van Stolk, C. (2017). Why Sleep Matters-The Economic Costs of Insufficient Sleep: A Cross-Country Comparative Analysis. Rand Health Q. 6, 11.

20. Streatfeild, J., Smith, J., Mansfield, D., Pezzullo, L., and Hillman, D. (2021). The social and economic cost of sleep disorders. Sleep 44, zsab132. 10.1093/sleep/zsab132.

21. Chaput, J.-P., Dutil, C., Featherstone, R., Ross, R., Giangregorio, L., Saunders, T.J., Janssen, I., Poitras, V.J., Kho, M.E., Ross-White, A., et al. (2020). Sleep duration and health in adults: an overview of systematic reviews. Appl. Physiol. Nutr. Metab. 45, S218–S231. 10.1139/apnm-2020-0034.

22. Grandner, M.A. (2017). Sleep, Health, and Society. Sleep Med. Clin. 12, 1–22. 10.1016/j.jsmc.2016.10.012.

23. Cappuccio, F.P., D’Elia, L., Strazzullo, P., and Miller, M.A. (2010). Sleep duration and all-cause mortality: a systematic review and meta-analysis of prospective studies. Sleep 33, 585–592. 10.1093/sleep/33.5.585.

24. Hirshkowitz, M., Whiton, K., Albert, S.M., Alessi, C., Bruni, O., DonCarlos, L., Hazen, N., Herman, J., Adams Hillard, P.J., Katz, E.S., et al. (2015). National Sleep Foundation’s updated sleep duration recommendations: final report. Sleep Health 1, 233–243. 10.1016/j.sleh.2015.10.004.

25. Valeggia, C.R., Burke, K.M., and Fernandez-Duque, E. (2010). Nutritional status and socioeconomic change among Toba and Wichí populations of the Argentinean Chaco. Econ. Hum. Biol. 8, 100–110. 10.1016/j.ehb.2009.11.001.

26. Fischer, D., Klerman, E.B., and Phillips, A.J.K. (2021). Measuring sleep regularity: theoretical properties and practical usage of existing metrics. Sleep 44, zsab103. 10.1093/sleep/zsab103.

27. Windred, D.P., Burns, A.C., Lane, J.M., Saxena, R., Rutter, M.K., Cain, S.W., and Phillips, A.J.K. (2024). Sleep regularity is a stronger predictor of mortality risk than sleep duration: A prospective cohort study. SLEEP 47, zsad253. 10.1093/sleep/zsad253.

28. Phillips, A.J.K., Clerx, W.M., O’Brien, C.S., Sano, A., Barger, L.K., Picard, R.W., Lockley, S.W., Klerman, E.B., and Czeisler, C.A. (2017). Irregular sleep/wake patterns are associated with poorer academic performance and delayed circadian and sleep/wake timing. Sci. Rep. 7, 3216. 10.1038/s41598-017-03171-4.

29. McKinnon, L., Samson, D.R., Nunn, C.L., Rowlands, A., Salvante, K.G., and Nepomnaschy, P.A. (2022). Technological infrastructure, sleep, and rest-activity patterns in a Kaqchikel Maya community. PLOS ONE 17, e0277416. 10.1371/journal.pone.0277416.

30. Samson, D.R. (2021). The Human Sleep Paradox: The Unexpected Sleeping Habits of Homo sapiens. Annu. Rev. Anthropol. 50, 259–274. 10.1146/annurev-anthro-010220-075523.

31. Iparraguirre, G. (2022). Cultural Rhythmics: Applied Anthropology and Global Development from Latin America (Emerald Publishing Limited) 10.1108/9781803828237.

32. R Core Team (2024). R: A Language and Environment for Statistical Computing (R Foundation for Statistical Computing).

33. Posit team (2024). RStudio: Integrated Development Environment for R (Posit Software, PBC).

34. Thieurmel, B., and Elmarhraoui, A. (2017). suncalc: Compute Sun Position, Sunlight Phases, Moon Position and Lunar Phase. 10.32614/CRAN.package.suncalc https://doi.org/10.32614/CRAN.package.suncalc.

35. Casiraghi, L., Spiousas, I., Dunster, G.P., McGlothlen, K., Fernández-Duque, E., Valeggia, C., and Iglesia, H.O. de la (2021). Moonstruck sleep: Synchronization of human sleep with the moon cycle under field conditions. Sci. Adv. 7, eabe0465. 10.1126/sciadv.abe0465.

36. Van Hees, V.T., and Migueles, J.H. (2013). GGIR: Raw Accelerometer Data Analysis. 10.32614/CRAN.package.GGIR https://doi.org/10.32614/CRAN.package.GGIR.

37. van Hees, V.T., Sabia, S., Jones, S.E., Wood, A.R., Anderson, K.N., Kivimäki, M., Frayling, T.M., Pack, A.I., Bucan, M., Trenell, M.I., et al. (2018). Estimating sleep parameters using an accelerometer without sleep diary. Sci. Rep. 8, 12975. 10.1038/s41598-018-31266-z.

38. Bland, J.M., and Altman, D.G. (1999). Measuring agreement in method comparison studies. Stat. Methods Med. Res. 8, 135–160. 10.1177/096228029900800204.

39. Leys, C., Ley, C., Klein, O., Bernard, P., and Licata, L. (2013). Detecting outliers: Do not use standard deviation around the mean, use absolute deviation around the median. J. Exp. Soc. Psychol. 49, 764–766. 10.1016/j.jesp.2013.03.013.

40. Delacre, M., and Klein, O. (2019). Routliers: Robust Outliers Detection. 10.32614/CRAN.package.Routliers https://doi.org/10.32614/CRAN.package.Routliers.

41. Bates, D., Maechler, M., Bolker, B., Walker, S., Christensen, R.H.B., Singmann, H., Dai, B., Scheipl, F., Grothendieck, G., Green, P., et al. (2025). lme4: Linear Mixed-Effects Models using “Eigen” and S4. Version 1.1-36.

42. Roenneberg, T., Kuehnle, T., Pramstaller, P.P., Ricken, J., Havel, M., Guth, A., and Merrow, M. (2004). A marker for the end of adolescence. Curr. Biol. 14, R1038–R1039. 10.1016/j.cub.2004.11.039.

43. Lüdecke, D., Makowski, D., Ben-Shachar, M.S., Patil, I., Højsgaard, S., and Wiernik, B.M. (2019). parameters: Processing of Model Parameters. 10.32614/CRAN.package.parameters https://doi.org/10.32614/CRAN.package.parameters.

44. Lüdecke, D., Makowski, D., Ben-Shachar, M.S., Patil, I., Waggoner, P., Wiernik, B.M., and Thériault, R. (2019). performance: Assessment of Regression Models Performance. 10.32614/CRAN.package.performance https://doi.org/10.32614/CRAN.package.performance.

45. Lenth, R.V. (2017). emmeans: Estimated Marginal Means, aka Least-Squares Means. 10.32614/CRAN.package.emmeans https://doi.org/10.32614/CRAN.package.emmeans.

46. de la Iglesia, H.O., Fernández-Duque, E., Golombek, D.A., Lanza, N., Duffy, J.F., Czeisler, C.A., and Valeggia, C.R. (2015). Access to Electric Light Is Associated with Shorter Sleep Duration in a Traditionally Hunter-Gatherer Community. J. Biol. Rhythms 30, 342–350. 10.1177/0748730415590702.

